# Differential temporal filtering in the fly optic lobe

**DOI:** 10.1101/2024.12.16.628613

**Authors:** Alexander Borst

## Abstract

Visual interneurons come in many different flavors, representing luminance changes at one location as ON or OFF signals with different dynamics, ranging from purely sustained to sharply transient responses. While the functional relevance of this representation for subsequent computations like direction-selective motion detection is well understood, the mechanisms by which these differences in dynamics arise are unclear.

Here, I study this question in the fly optic lobe. Taking advantage of the known connectome I simulate a network of five adjacent optical columns each comprising 65 different cell types. Each neuron is modeled as an electrically compact single compartment, conductance-based element that receives input from other neurons within its column and from its neighboring columns according to the intra- and inter-columnar connectivity matrix. The sign of the input is determined according to the known transmitter type of the presynaptic neuron and the receptor on the postsynaptic side. In addition, some of the neurons are given voltage-dependent conductances known from the fly transcriptome. As free parameters, each neuron has an input and an output gain, applied to all its input and output synapses, respectively. The parameters are adjusted such that the spatio-temporal receptive field properties of 13 out of the 65 simulated neurons match the experimentally determined ones as closely as possible.

Despite the fact that all neurons have identical leak conductance and membrane capacitance, this procedure leads to a surprisingly good fit to the data, where specific neurons respond transiently while others respond in a sustained way to luminance changes. This fit critically depends on the presence of an H-current in some of the first-order interneurons, i.e., lamina cells L1 and L2: turning off the H-current eliminates the transient response nature of many neurons leaving only sustained responses in all of the examined interneurons. I conclude that the diverse dynamic response behavior of the columnar neurons in the fly optic lobe starts in the lamina and is created by their different intrinsic membrane properties. I predict that eliminating the hyperpolarization-activated current by RNAi should strongly affect the dynamics of many medulla neurons and, consequently, also higher-order functions depending on them like direction-selectivity in T4 and T5 neurons.

**Author summary:** Visual interneurons come in many different flavors, representing luminance changes at one location as ON or OFF signals and with different dynamics, ranging from purely sustained to sharply transient. While the functional relevance of this representation for subsequent computations, like direction-selective motion detection, is well understood, the mechanism by which these differences in dynamics arise is unclear. Here, I study this question by using the connectome of the fly optic lobe and simulating the network of interneurons in a biophysically plausible way. Adjusting the input and the output gain of each neuron such that a subset of neurons (those where experimental data exist) matches the response kinetics of their real counterparts, I identify a particular voltage-gated ion channel present in some of the first-order interneurons as being critical for the transient response behavior of postsynaptic neurons. This study, therefore, predicts that eliminating this current from the circuit should strongly affect the response kinetics in downstream circuit elements and destruct the computation of direction selectivity.

## Introduction

Luminance information represented by the activity pattern of the photoreceptors is processed in subsequent neuropil layers in many different parallel channels. As a result, visual interneurons come in a variety of different flavors along various dimensions: Cells can be classified as ON or OFF cells depending on their contrast polarity preference, cells can differ in their receptive field size with more or less pronounced antagonistic surrounds, and cells can be called sustained or transient, depending on their response time course. A telling example comes from the mouse retina where bipolar cells projecting to different adjacent layers in the inner retina smoothly vary from OFF sustained to OFF transient and ON transient to ON sustained in a systematic way [1]. In a similar way, visual interneurons in the fly optic lobe are either ON or OFF cells [2,3] and respond to luminance changes in a transient or sustained way (Fig 1, modified from [4]; see also [3,5,6]). Differential temporal filtering of retinal inputs is important for subsequent computations like visual motion detection which requires an interaction of a delayed signal from one retinal location with an un-delayed signal from an adjacent retinal location [7-9]. However, the mechanism by which such differences in temporal properties arise is unclear, in the vertebrate retina as well as in the fly optic lobe.

**Figure 1.**
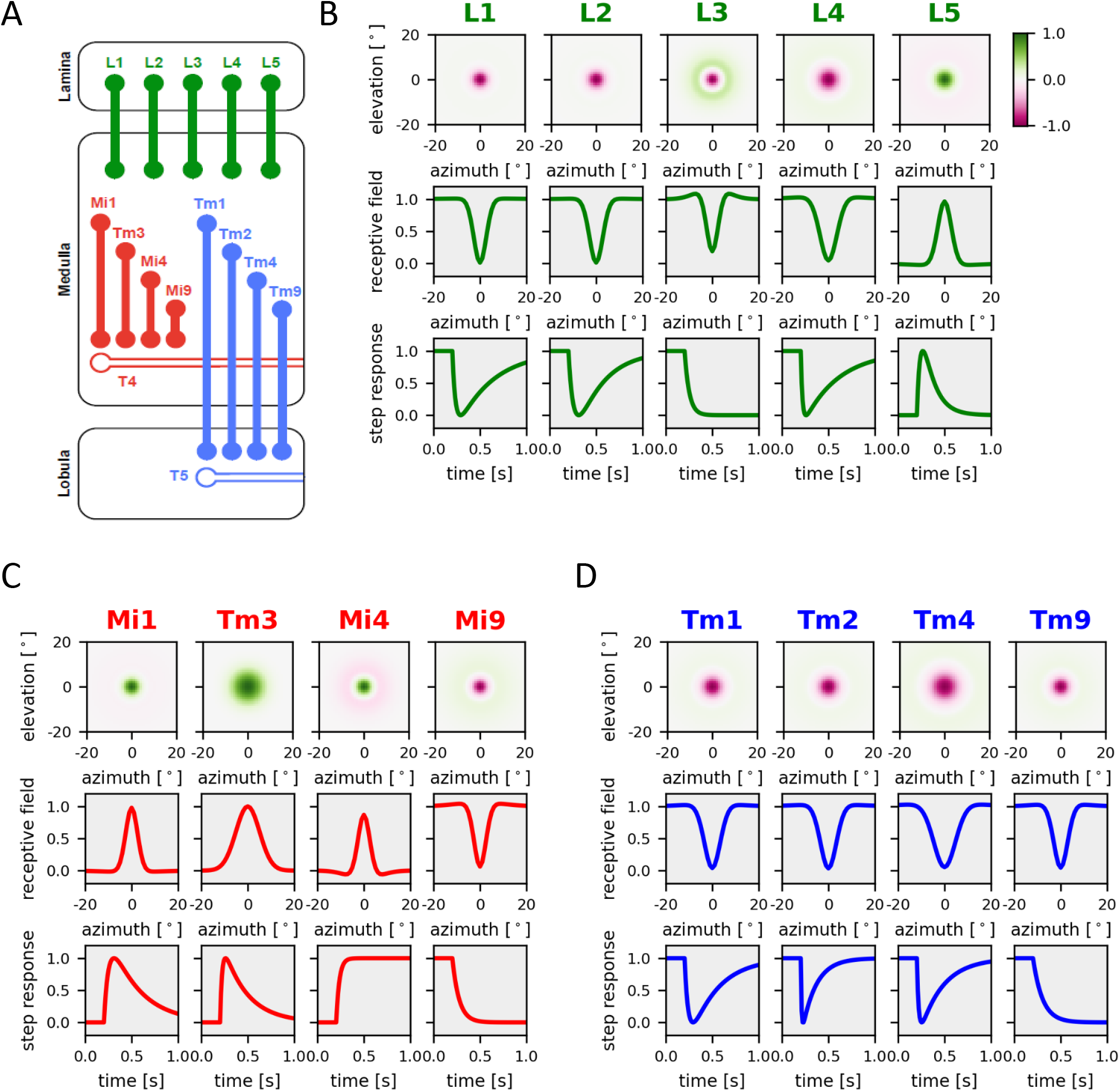
Linear receptive fields of columnar neurons obtained by reverse correlation from calcium responses to stochastic stimuli in tethered fruit flies. (**A**) Schematic of the circuit including all columnar neurons presynaptic of T4 and T5, which have been characterized (graphics from [25]). ON-pathway neurons are shown in red, OFF-pathway neurons in blue, lamina neurons in green. (**B**) Results for lamina columnar neurons L1–5. *Top*: False-color plot of the linear spatial receptive fields from a Difference-of-Gaussian model fit to the original data. *Middle*: 1D spatial profile of the receptive field. *Bottom:* Step responses of the cells (data replotted from [25]). (**C**) Same as in C for columnar elements of the ON-pathway, Mi1, Tm3, Mi4 and Mi9. (**D**) Same as in C for columnar elements of the OFF-pathway, Tm1, Tm2, Tm4 and Tm9 (data in C and D replotted [24,25]).

In principle, the temporal dynamics of a neuron’s response to a sensory stimulus are shaped by its intrinsic membrane properties as well as by its input signal. Intrinsic membrane properties comprise passive properties, i.e., membrane leak conductance and membrane capacitance, and active properties, i.e., voltage-dependent conductances for various ions. The input signal may consist of the primary sensory receptor output signal, in case it is directly postsynaptic to the sensory receptor cell, as well as the output signals of all other neurons it is connected to it. In the latter case, one may distinguish between a pure feed-forward chain and recurrent feed-back connectivity. In general, thus, the neural response time-constants are a combination of the intrinsic time-constants and the connectivity which can lead to response time-constants exceeding by far the ones of any individual neuron in the network [10,11]. Given the wealth of information available for neurons in the fly optic lobe, which includes the connectome [12], the transcriptome [13] and physiological recording data from either calcium-imaging or patch-clamp experiments, I decided to investigate the mechanistic origin of differential temporal filtering by modeling the network of optic-lobe neurons with biophysically plausible, conductance-based elements. Adjusting the input and the output gain of each neuron such that a subset of neurons (those where experimental data exist) matches the response kinetics of their real counterparts, I identify a particular voltage-gated ion channel present in some of the first-order interneurons as being critical for the transient response behavior of postsynaptic neurons. I conclude that transient behavior of most second-order interneurons is not created by feed-back loops but rather inherited from first-order interneurons.

## Results

In order to capture the spatial and the temporal properties of columnar neurons in the fly visual system, I simulated the neural activity in five adjacent columns arranged in a linear way (Fig 2A). Each column contained 65 cell types, each with identical properties across all columns. The connectivity within each column (Fig 2B) was set according to connectomic data [14], with the strength of each connection corresponding to the number of synapses and the sign based on the known transmitter phenotype of each cell and its postsynaptic receptor. The connectivity between neighboring columns (Fig 2C) was constructed in the same way, except for L4 cells, whose branches in neighboring columns were assumed to be electrically isolated. This resulted in a network of 325 neurons with an overall connectivity combining the intra- and inter-columnar connections (Fig 2D).

**Figure 2.**
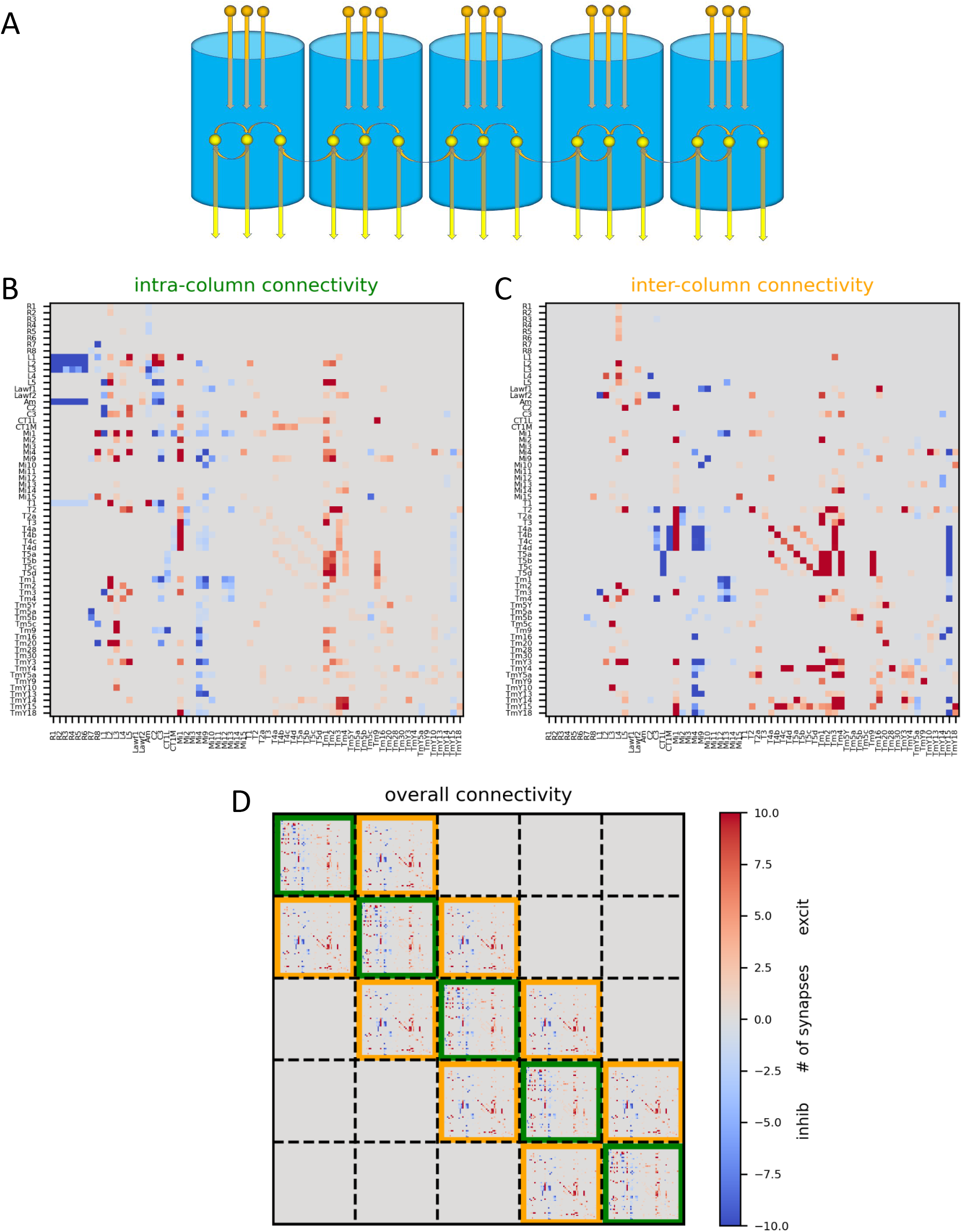
Layout of network model. (**A**) Schematic of the circuit. Five adjacent columns, arranged in a linear way, were simulated. (**B**) Intra-columnar connectivity matrix containing the synaptic weights between all 65 columnar elements. (**C**) Same as B, but for inter-columnar connectivity. (**D**) Resulting overall connectivity matrix for all 325 network elements. Data from [14], split into intra- and inter-columnar connectivity by Janna Lappalainen.

**Figure 3.**
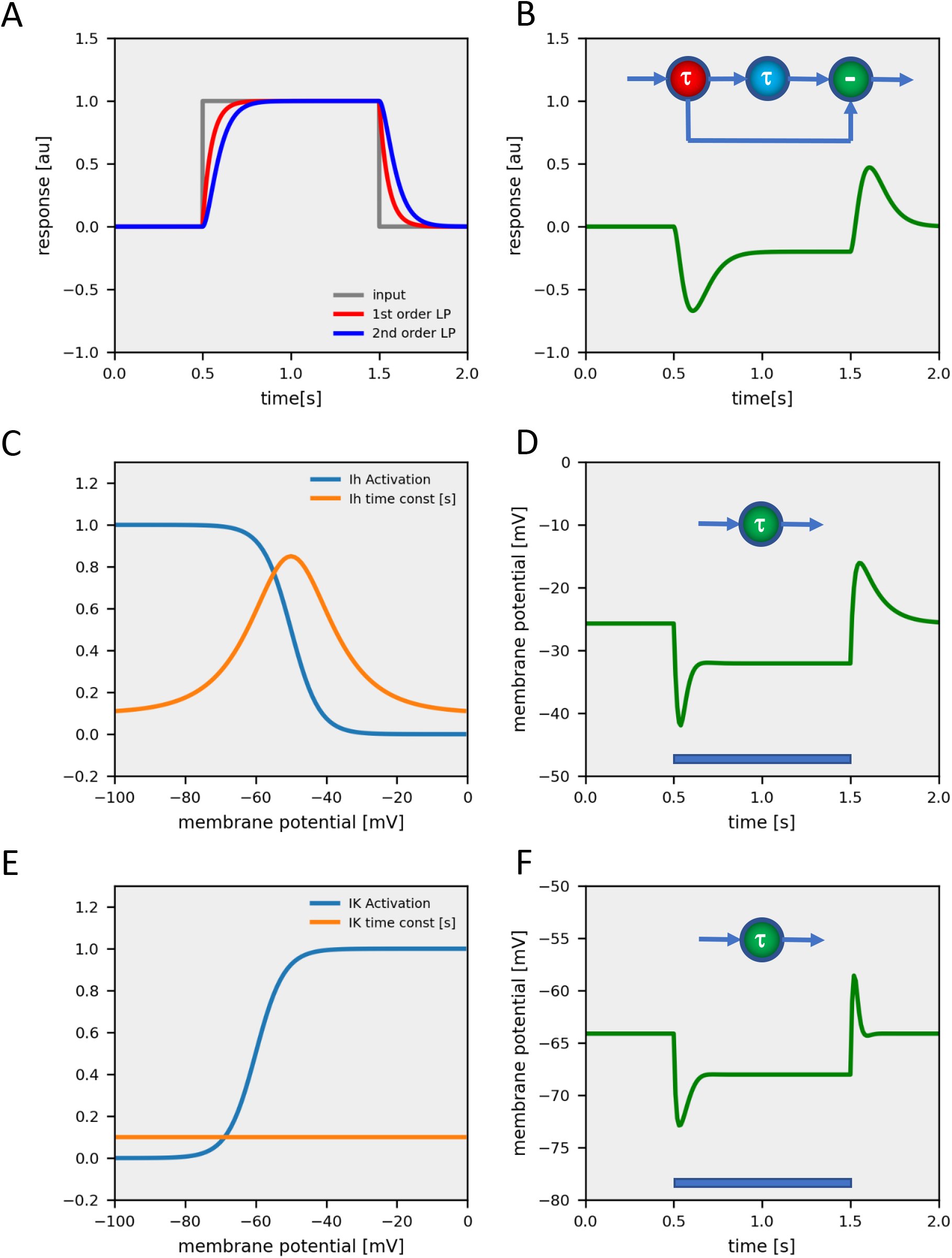
Different ways to produce transient responses. (**A**) Responses of a 1^st^ (in red) and of a 2^nd^ order lowpass filter (in blue) to a pulse of 1 s duration. (**B**) The unbalanced difference of 1^st^ and 2^nd^ order signals leads to a transient hyperpolarization response. (**C**) Activation characteristic (in blue) and voltage-dependent time-constant (in orange) of a H-current. (**D**) Response of a model neuron with an H-current to a hyperpolarizing current injection. (**E**) Activation characteristic (in blue) and time-constant (in orange) of a non-inactivating K-current. (**F**) Response of a model neuron with a K-current to a hyperpolarizing current injection.

I modeled each neuron as a single-compartment, conductance-based, graded-potential element (for detail, see Material and Methods). Each neuron received excitatory and inhibitory input according to the overall connectivity matrix. Importantly, each neuron type was given two free parameters, an input and an output gain. By applying the input gain of a neuron to all its presynaptic input neurons and the output gain uniformly to all its postsynaptic downstream neurons, the relative strength of connectivity as set by the connectivity matrix was preserved. All neurons were given the same capacitance and leak conductance, and, thus, had uniform intrinsic passive membrane properties. Lamina cells L1-3 are the first neurons postsynaptic to photoreceptors R1-6, and, therefore, should be important for shaping the response properties of downstream medulla neurons. L1 and L2 are known to respond to a light pulse by a transient hyperpolarization with a typical ‘sag’ in the neuron’s membrane potential, followed by a rebound excitation after the light has been turned off [15,16]. Such a response can be caused by two different mechanisms: (1) Low-pass filtering the light response with two different time-constants and taking the difference between the two outputs (Fig 2A,B) or by (2) intrinsic, active membrane properties of the lamina cells (Fig 2C-F). In the latter case, the current leading to the response sag could be a depolarizing current activated by hyperpolarization such as the H-current (Fig 2C,D), or the deactivation of a hyperpolarizing current, such as a non-inactivating K-current (Fig 2E,F).

Transcriptomic data indicated a high expression level of the gene coding for an HCN-channel (FlyBase ID: FBgn0263397), responsible for an H-current, in lamina cells L1, L2 and L5 [13]. I, therefore, incorporated such a current in lamina cells in some of the simulations. With each neuron type having an input and an output gain, the total number of free parameters in the model amounted to 130. If the model incorporated an H-current, 5 additional parameters were used to describe the maximum conductances in the five lamina cells and 3 parameters to characterize the voltage-dependence of the H-current activation curve and time-constant (for details, see Material and Methods).

To determine the spatio-temporal receptive field properties of the model neurons, a depolarizing current step was injected into photoreceptors R1 to R8 within the central column, equivalent to a light pulse delivered locally. Its spread into the other neighboring columns was used to determine the spatial receptive field of each neuron type. To account for the fact that experimental data were derived from calcium imaging, the voltage responses of the model neurons were finally fed through a 1^st^-order low-pass filter with 50 ms time-constant. To fit the model responses to the data, an overall error or cost of the model was defined as the squared difference between data and model responses divided by the data power. Optimal parameter sets were found by minimizing the cost function by gradient descent starting from a random parameter set.

In a first round of simulations, I investigated to what extent all the different dynamics of fly visual interneurons can be accounted for using identical, uniform, passive membrane properties of all model neurons. Thus, the only way to achieve different response properties is by the network connectivity. The resulting receptive field properties from such simulations overall matched the experimental data in quite amazing detail, leading to a final cost of about 7 % (Fig 4). Importantly, the pronounced antagonistic surround of lamina neuron L3 and medulla neuron Mi4 was well captured by the model, indicating that inter-columnar connectivities can account for their spatial receptive field profiles. Furthermore, and of particular interest in the current context, the transient responses of lamina neurons L1, L2 and L4, as well as of medullar neurons Mi1, Tm3, Tm1, Tm2 and Tm4 were replicated by the model, together with the sustained responses of L3, Mi4, Mi9 and Tm9. However, unlike its biological counterpart, lamina neuron L5 responded in a sustained way. Furthermore, its response amplitude, as well as the one of medulla neuron Tm4, fell short of the respective measured amplitudes. As a final note, many responses showed ringing, indicating some sort of instability within feed-back loops.

**Figure 4.**
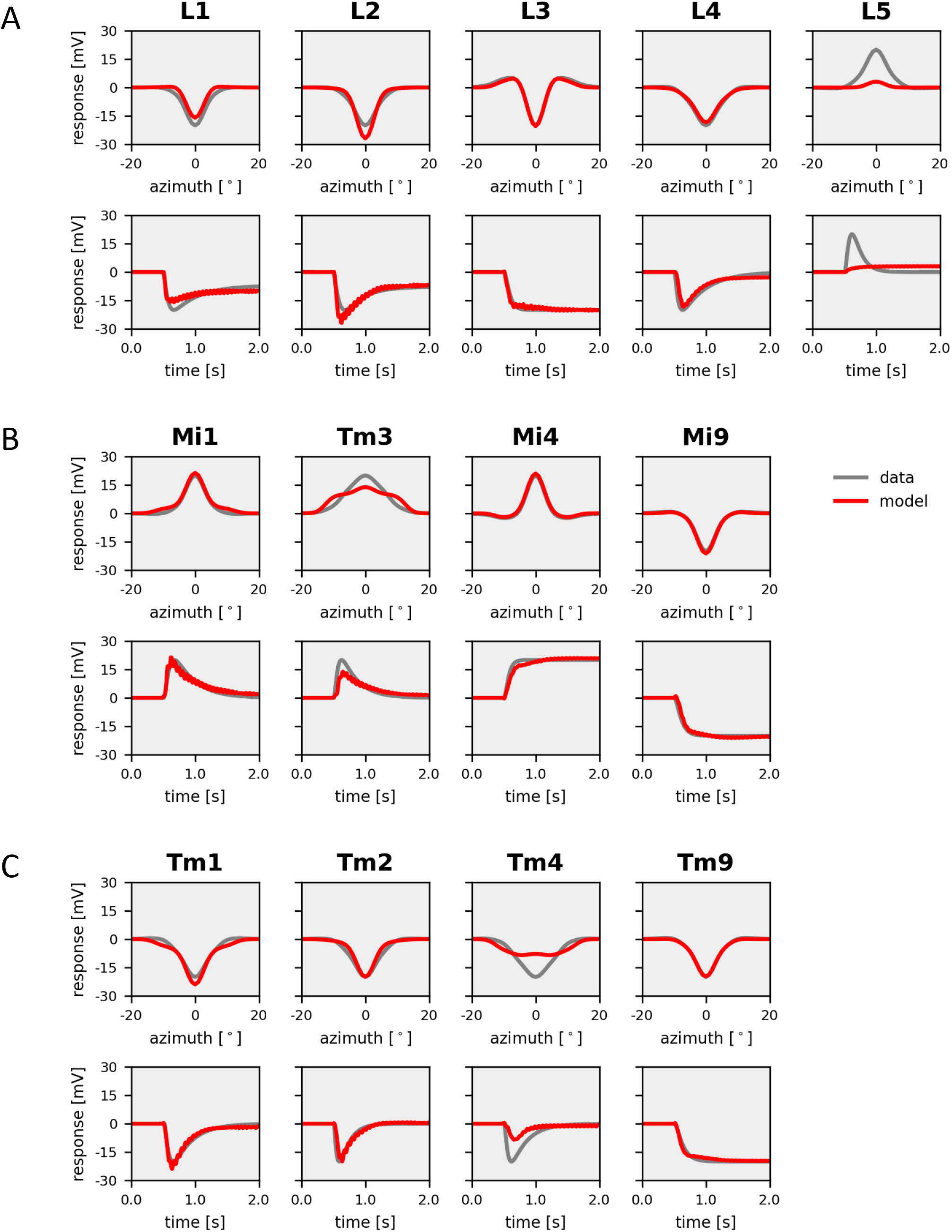
Model simulation with electrically passive elements. Responses of model neurons (in red) with parameters optimized to fit experimental data (in gray). (**A**) *Top*: Spatial profiles of the receptive field of the five lamina cells L1-L5. *Bottom*: Temporal step response of the same neurons. (**B**) Same as A, but for columnar elements of the ON-pathway, Mi1, Tm3, Mi4 and Mi9. (**C**) Same as A, but for columnar elements of the OFF-pathway, Tm1, Tm2, Tm4 and Tm9.

Next, I inserted an H-current into lamina neurons and minimized the cost, again starting from a random parameter set. This time, the model had 5 additional parameters determining the maximum conductance in each of the 5 lamina neurons and 3 parameters describing the voltage characteristics of the H-current, shared for all 5 lamina neurons. The resulting receptive field properties of the 13 selected neuron types were in most cases almost indistinguishable from the experimental data, with a final cost of only 3 % (Fig 5). As in the data, the spatial profiles of the receptive fields of lamina cell L3 and medulla neuron Mi4 had antagonistic surrounds, and that of medulla cell Tm3 was extending over several columns. The only significant deviation was seen in medulla neurons Tm3 and Tm4 where the model responses had a smaller amplitude than the real neurons. All other cells faithfully matched the data in their spatial profile as well as in their time-course. Furthermore, ringing was almost completely absent in the responses. Most importantly, the distinction between transient and sustained cells in the model was as clear as in the data.

**Figure 5.**
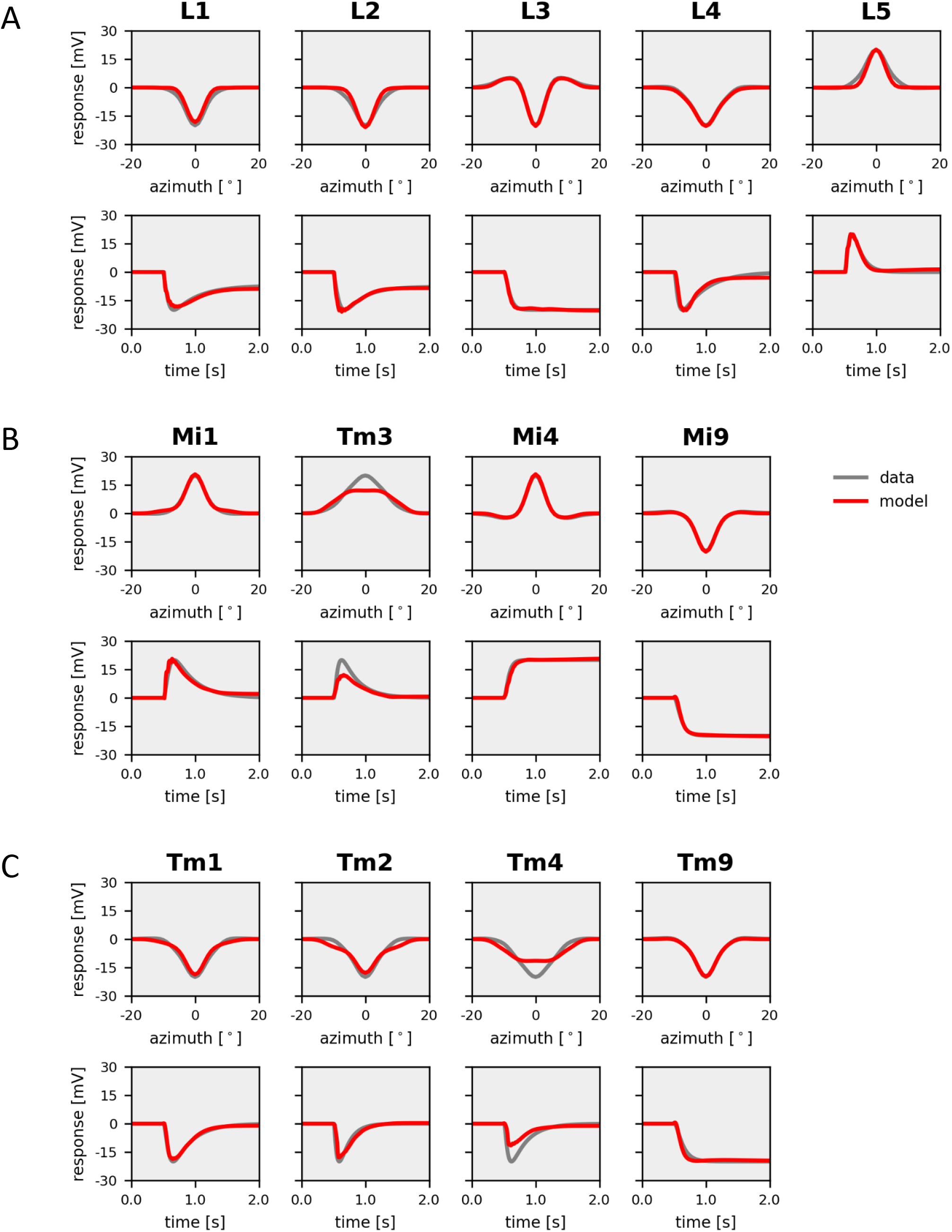
Model simulation with H-current in L1, L2 and L5. Responses of model neurons (in red) with parameters optimized to fit experimental data (in gray). (**A**) *Top*: Spatial profiles of the receptive field of the five lamina cells L1-L5. *Bottom*: Temporal step response of the same neurons. (**B**) Same as A, but for columnar elements of the ON-pathway, Mi1, Tm3, Mi4 and Mi9. (**C**) Same as A, but for columnar elements of the OFF-pathway, Tm1, Tm2, Tm4 and Tm9.

Next, I investigated to what extent the dynamic properties of this network still depend on the circuit connectivity or, alternatively, now are largely dependent on the intrinsic, transient membrane properties of lamina neurons L1, L2, L4 and L5. To this end, I switched off the H-current in the above network after optimizing the parameters with an H-current. This procedure led to a dramatic break-down of the match between model and data, quantified by a cost of more than 100 % (Fig 6). Importantly, all model neurons now responded in a sustained way, i.e., have lost their transient nature.

**Figure 6.**
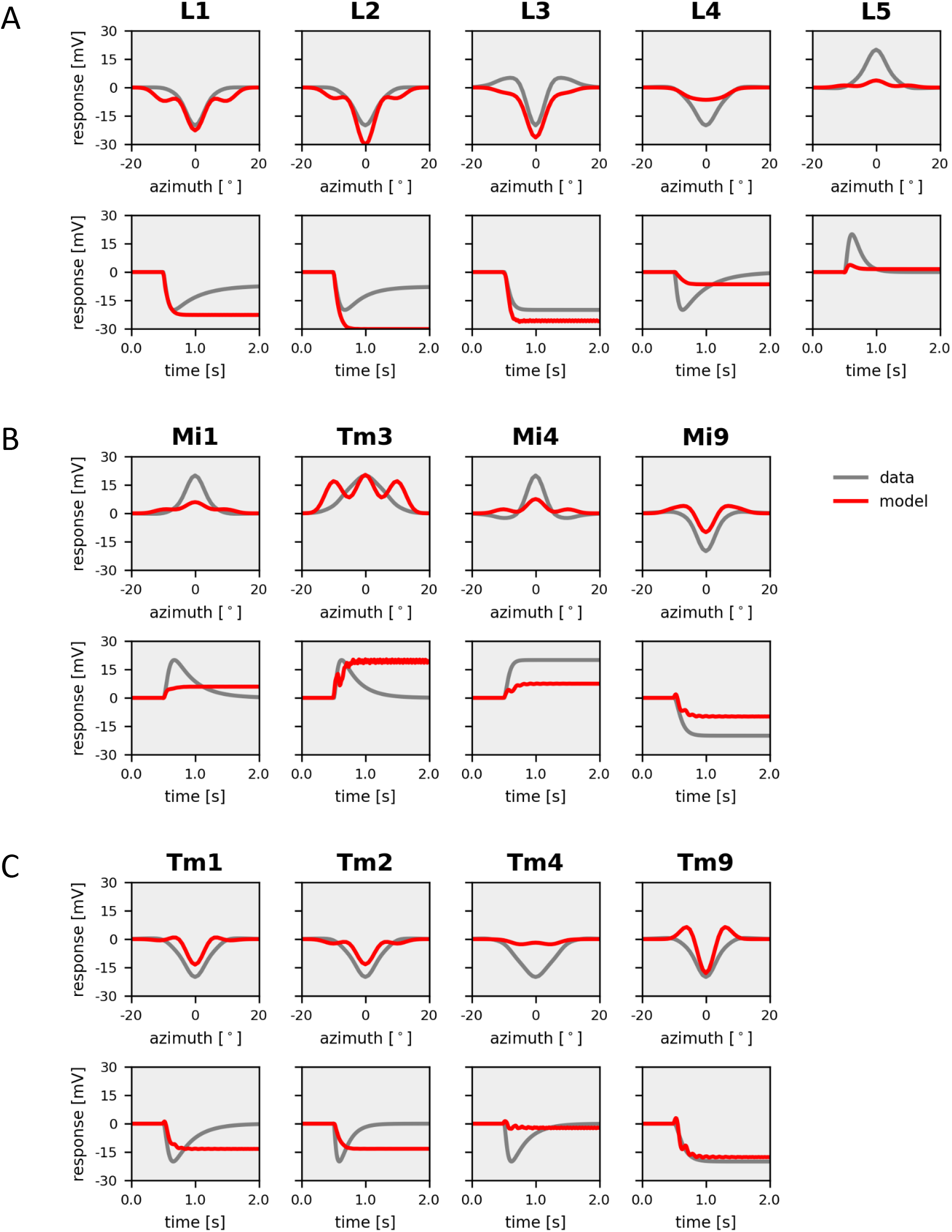
Model simulation with H-current shut off in L1, L2 and L5. Responses of model neurons (in red) after parameters optimized with H-current to fit experimental data (in gray). (**A**) *Top*: Spatial profiles of the receptive field of the five lamina cells L1-L5. *Bottom*: Temporal step response of the same neurons. (**B**) Same as A, but for columnar elements of the ON-pathway, Mi1, Tm3, Mi4 and Mi9. (**C**) Same as A, but for columnar elements of the OFF-pathway, Tm1, Tm2, Tm4 and Tm9.

I finally tested the consistency of the above conclusions by comparing the parameter sets of the 10 best models obtained from 20 optimization runs in both scenarios, i.e., without and with an H-current. Without an H-current, final model costs ranged from 7% to about 11% (Fig 7A), while costs of the 10 best models with an H-current all were found below 4% (Fig 7B). This means that introduction of an H-current led to a cost reduction of about 50%. In each model category, parameters were highly correlated (Fig 7C,D) indicating that different optimization runs converged on a similar parameter set each time. However, these parameter sets were significantly different for models without and with an H-current (Fig 7E,F), with input and output gains being in general smaller for models without an H-current compared to models with an H-current. Interestingly, despite the fact that all five lamina cells were given an H-current, optimization consistently converged on models where only lamina cells L1 and L2 had a large maximum H-current conductance, a small one in lamina cells L4 and L5, and zero in lamina cell L3, which is in approximate agreement with the transcriptomic data of Davis et al, 2020 [13].

**Figure 7.**
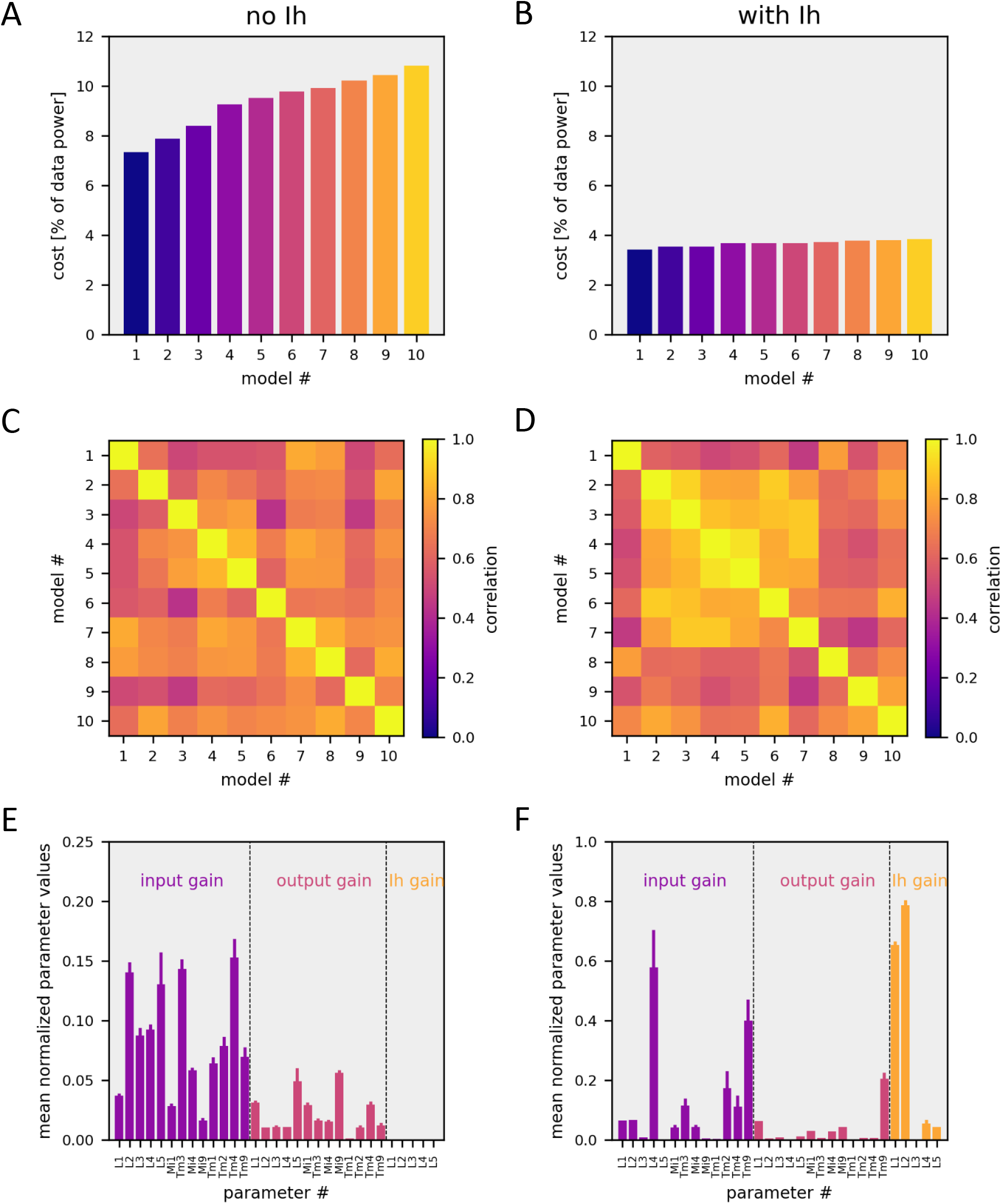
Model Evaluation. Shown are the 10 best models from 20 optimization runs. (**A**,**B**) Remaining cost values of models without (A) and with (B) an H-current. (**C**,**D**) Correlation matrix showing the Pearson Correlation Coefficient of all parameters between each of the 10 models, without (C) and with (D) an H-current. (**E**,**F**) Average values +-variance of selected model parameters from the 10 best models without (E) and with (F) an H-current.

## Discussion

The model simulations presented above show that, in principle, it is possible to replicate the response dynamics of sustained and transient lamina and medulla neurons by using identical membrane time-constants in all model neurons (Fig 4). In this case, all differences are due to the connectivity within the network. However, inserting an H-current into lamina neurons improved the fit to the data significantly (Fig 5). Turning off the H-current then led all neurons to respond in a sustained way (Fig 6) emphasizing the critical role of this current in lamina cells for the overall response dynamics of the circuit elements in the fly optic lobe.

The transient hyperpolarization of lamina monopolar cells in response to light stimuli have been known for long [15,16]. It is caused by a histamine release from fly photoreceptors opening a chloride conductance [17]. Assuming a rather constant transmitter release from photoreceptors during light stimulation, the transient nature of the hyperpolarization in the postsynaptic lamina cells could come about by an inactivation of the histamine receptor or by a subsequent depolarizing current. Such a depolarizing current can be caused extrinsically, i.e. through synaptic connectivity with other interneurons, or by an intrinsic current such as the hyperpolarization-activated H-current (Fig 3C,D). Interestingly, the gene encoding for the HCN-channel underlying an H-current is transcribed in 11 isoforms, one of which was found to be highly expressed in lamina cells L1 and L5, at an intermediate level in L2 and L4, and zero in L3 [13]. Given the strong electrical coupling between L1 and L2 [2], this expression pattern is in principle in agreement with the transient response behavior of L1 and L2 cells on the one, and the sustained responses of L3 on the other side. However, none of the HCN-isoforms has so far been characterized in electrophysiological experiments, leaving their specific properties in the dark and open to parameter adjustment. The optimal parameters found in the simulations above agree qualitatively with the ones determined for H-currents of thalamic neurons in vertebrates [18]. Interestingly, maximal conductance of this H-current, set as a free parameter for all 5 lamina cells, was consistently found to be highest in L1, L2, small in L4 and L5, and zero for L3 in all simulation runs.

K-channels are another way to change the membrane conductance and, hence, can influence the membrane time-constant. Indeed, various K-channels have been shown to affect the response time-course of *Drosophila* neurons [19,20]. Specifically, two A-type K-channels -*Shaker* and *Shal* – are expressed in L2 and L3 cells, and, when blocked, either pharmacologically or genetically, lead to somewhat slower responses [20]. However, such inactivating K-channels can only affect the rebound depolarization seen in L1 and L2, but not the sag in their hyperpolarizing response to light [21]. For the latter, a non-inactivating K-channel would be required with the resting membrane potential of the neuron rather hyperpolarized (Fig 3E,F). Lamina neurons L1 and L2, however, have a resting potential of around −38 mV [15,21].

The optic lobe network has recently been modeled by Lappalainen and colleagues [22]. Pretty good matches were found in this study between the various model and real neurons of the fly optic lobe. The simulation study presented here differs from that of Lappalainen et al. in the following four aspects: (1) Lappalainen et al (2024) trained the network to perform a certain task, i.e., to extract the optic flow from a movie. While optimizing the network in this respect, they looked at the receptive fields of the various visual interneurons only as a corollary. In contrast, I used the match between the modeled and measured responses of selected visual interneurons as a cost for parameter optimization. (2) Lappalainen et al (2024) used recti-linear model elements, whereas the neurons in my simulation were all conductance-based. (3) Lappalainen et al (2024) had the synaptic strength of each non-zero connection of the connectome as a free parameter, whereas I used a uniform input and output gain of each neuron applied to all its input and output synapses according to the weights defined in the connectome. This not only led to fewer free parameters in my study (138 here vs 734 in Lappalainen et al, 2024), it also led to more stringent confinement by the connectomic data. (4) Most importantly, in the context of the question investigated in my study, Lappalainen et al (2024) had the time-constant of each individual model element as a free parameter, whereas all neurons in my study had identical leak conductance and membrane capacitance: when isolated from the network, this would result in identical membrane time-constants.

The main result of the network simulation presented above is that the difference in intrinsic membrane properties of the first order interneurons, i.e., L1-L3, is most critical for the dynamics of all subsequent medulla neurons. Specifically, the H-current expressed in lamina cells L1 and L2 but not in L3 determines the transient vs sustained nature of medulla neurons providing input to motion-sensitive neurons T4 in the ON- and T5 in the OFF-pathway. Since the different dynamics of input neurons is a prerequisite for the computation of the motion direction [7,9,23], this study further predicts that knocking down the transcript of the gene encoding for the HCN-channel by RNAi should not only alter response dynamics of columnar medulla neurons, but also strongly impair direction-selectivity in T4 and T5 cells.

## Material and Methods

### Model

Each neuron *i* was modeled as an electrically compact single-compartment, conductance-based graded element, using the following, first-order differential equation:

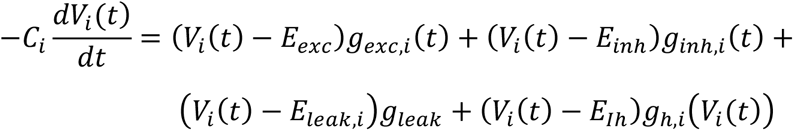

This equation states that the sum of five different currents equals zero. The first one is the capacitive current 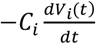. The next two terms, (*V*_*i*_ (*t*) − *E*_*exc*_)*g*_*exc*,*i*_ (*t*), and (*V*_*i*_ (*t*) − *E*_*inh*_)*g*_*inh*,*i*_ (*t*) describe the excitatory and inhibitory synaptic currents. For that, the overall connectivity matrix **M** was split into an excitatory and an inhibitory matrix, **M**_exc_ and **M**_inh_, respectively, containing non-negative entries only. The total excitatory and inhibitory conductance of each neuron i was then calculated as the sum over all its inputs, weighted by the synaptic strength as derived from the two connectivity matrices:

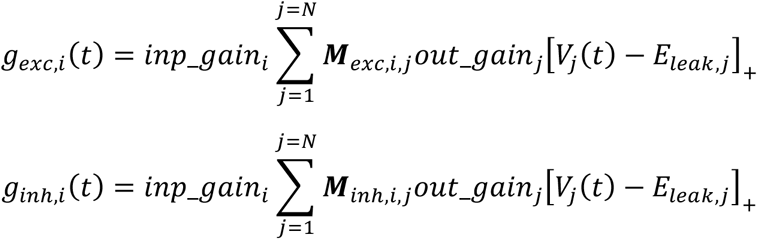

In these calculations, the output of each neuron *j* was rectified with respect to its leak potential *E*_leak_. *E*_leak_ was set to −50 mV for all neurons in the circuit, except for lamina cells L1-L3 where *E*_leak_ was set to −20 mV. As is seen from the above equations, two parameters come into play here: the input gain of neuron i and the output gain of neuron j. By applying the output gain of a neuron uniformly to all its postsynaptic downstream neurons, and the input gain to all its presynaptic input neurons, the relative strength of connectivity as set by the number of synapses determined in the EM, was preserved. The next term is (*V*_*i*_ (*t*) – *E*_*leak*,*i*_)*g*_*leak*_, which describes the ohmic leak current. Importantly, all neurons were given the same capacitance and leak conductance, and, thus had uniform intrinsic passive membrane properties. The last term in membrane equation, (*V*_*i*_(*t*) − *E*_*Ih*_)*g*_*h*,*i*_(*V*_*i*_(*t*)), describes an active, i.e., voltage-dependent, H-current. The gene encoding the underlying ion channel HCN is highly expressed in lamina cells L1, L2 and L5 [13]. In part of the model simulations, this current was inserted into the five lamina cells, in other simulations it was not. I modeled the H-current by two functions, its steady-state activation characteristic *g*_*h*,∞_, and its voltage-dependent time-constant [18]. The first was defined as a logistic function, involving the parameters maximum conductance *g*_*max*_, the voltage of half-maximum steady-state activation *V*_*mid*_, and the slope of the curve at this point, i.e., *slope*:

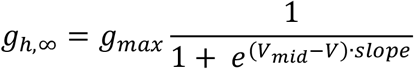

The voltage-dependent time-constant was defined by the following equation, involving one parameter, *τ*_*midV*_ :

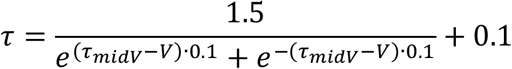

The conductance of the H-current in neuron i at each time was obtained from numerical integration of the following 1^st^-oder differential equation at a temporal resolution of 10 ms:

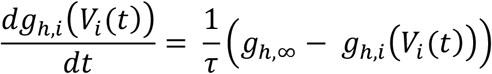

With each neuron having an input and an output gain, the total number of free parameters, thus, amounted to 130. If the model incorporated an H-current, 8 additional parameters (5 *g*_*max*_, *V*_*mid*_, *slope, τ*_*midV*_) were used, so total number of free parameters increased to 138.

To determine the spatio-temporal receptive field properties of the model neurons, a depolarizing current step was injected in photoreceptors R1 to R8 within the central column. Its spread into the other neighboring columns was used to determine the spatial receptive field of each neuron type in the central column. The time course of each neuron’s response described its temporal dynamics over 2 seconds at a temporal resolution of 10 ms, i.e., in T = 200 time steps. To account for the fact that experimental data were derived from calcium imaging, the voltage responses of the model neurons were finally fed through a 1^st^-order low-pass filter with 50 ms time-constant. Model responses were evaluated with respect to the membrane potential at stimulus onset, which was at t = 0.5 s. Prior to that, model responses were set to zero.

### Data

Data were taken from two previous studies [24,25]. There, the impulse responses of 13 neuronal cell types of the fly optic lobe were determined by calcium imaging, using white noise stimuli and reverse correlation. These data represented normalized impulse responses which were integrated over time to result in step responses and contained no information about membrane voltage. In order to compare them with simulations producing membrane potential output, I arbitrarily set the maximum of all data to 20 mV response magnitude. To account for the response characteristics of lamina cells L1 and L2 as seen in electrophysiological recordings (e.g. [16]), a DC level of 40% of the peak value was added to their step responses.

### Cost function

Denoting the free parameters as a vector **z**, the model responses of N = 13 out of the 65 neuron types were compared to the experimental data set (Fig 1) by the following cost function:

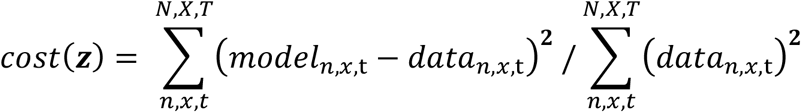

The cost function expresses the deviation of the model from the data responses as a fraction of the data power and, therefore, allows for a meaningful interpretation.

### Parameter Optimization

Optimal parameter sets were found by minimizing the cost function by gradient descent using the Adam optimizer of the PyTorch library. Starting from a random parameter set, the optimizer was applied in three consecutive rounds with 10,000 steps each, decreasing the learning rate from 0.1 to 0.01 to 0.001. This was repeated 20 times, resulting in 20 different models with an H-current, and 20 models without an H-current. The model with the smallest final cost value is shown in Fig 4 (optimized without an H-current) and Fig 5 (optimized with an H-current). The CPU time of one optimization run with 3 times 10 000 steps was about 25 minutes.

## Acknowledgements

I thank Janne Lappalainen for providing the intra- and inter-columnar connectivity matrices, Franz Rieger for help with PyTorch, and Juergen Haag, Georg Ammer and Lukas Groschner for comments on the manuscript. This work was supported by the Max Planck Society.

## Data and Code Availability

All data and python code are available on github.

